# plasmidSPAdes: Assembling Plasmids from Whole Genome Sequencing Data

**DOI:** 10.1101/048942

**Authors:** Dmitry Antipov, Nolan Hartwick, Max Shen, Mikhail Raiko, Alla Lapidus, Pavel A. Pevzner

## Abstract

**Motivation:** Plasmids are stably maintained extra-chromosomal genetic elements that replicate independently from the host cell’s chromosomes. Although plasmids harbor biomedically important genes, (such as genes involved in virulence and antibiotics resistance), there is a shortage of specialized software tools for extracting and assembling plasmid data from whole genome sequencing projects.

**Results:** We present the plasmidSPAdes algorithm and software tool for assembling plasmids from whole genome sequencing data and benchmark its performance on a diverse set of bacterial genomes.

**Availability and implementation:** PLASMIDSPADES is publicly available at http://spades.bioinf.spbau.ru/plasmidSPAdes/

## 1 INTRODUCTION

Plasmids are common in Bacteria and Archaea, but have been detected in Eukaryotes as well (Gunge *et al.*, 1982). The cells often have multiple plasmids of varying sizes existing together in different numbers of copies per cell. Plasmids are important genetic engineering tools and the vectors of horizontal gene transfer that may harbor genes involved in virulence and antibiotic resistance. Thus, studies of plasmids are important for understanding the evolution of these traits and for tracing the proliferation of drug-resistant bacteria.

Since plasmids are difficult to study using Whole Genome Sequencing (WGS) data, biologists often use special biochemical methods for extracting and isolating plasmid molecules for further *plasmid sequencing* (Williams *et al.*, 2006; Kav *et al.*, 2012). In the case of WGS, when a genome of a bacterial species is assembled, its plasmids often remain unidentified. Obtaining information about plasmids from thousands of genome sequencing projects (without preliminary plasmid isolation) is difficult since it is not clear which contigs in the genome assembly have arisen from plasmids.

Since the proliferation of plasmids carrying antimicrobial resistance and virulence genes leads to the proliferation of drug resistant-bacterial strains, it is important to understand the epidemiology of plasmids and to develop plasmid typing systems. Carattoli *et al.* (2014) developed PlasmidFinder software for detecting and classifying variants of known plasmids based on their similarity with plasmids present in plasmid databases. However, PlasmidFinder is unable to identify novel plasmids that have no significant similarities to known plasmids.

Lanza *et al.* (2014) developed the PLAsmid Constellation NET-work (PLACNET) tool for assembling plasmids from WGS data and applied it for analyzing plasmid diversity and adaptation (de Toro *et al.*, 2014) PLACNET uses three types of information to identify plasmids: (i) information about scaffold links and coverage in the WGS assembly, (ii) comparison to reference plasmid sequences, and (iii) plasmid-diagnostic sequence features such as replication initiator proteins. PLACNET combines these three types of data and outputs a network that needs to be further pruned by expert analysis to eliminate confounding data.

While combining all three types of data for plasmid sequencing is important, the focus of this paper is only on using WGS assembly for plasmid reconstruction. We argue that while the analysis of scaffolds in Lanza *et al.* (2014) is important, there is a wealth of additional information about plasmids encoded in the structure of the *de Bruijn graph* (constructed from *k*-mers in reads) that Lanza *et al.* (2014) do not consider. Recently, Rozov *et al.* (2015) demonstrated how to use the de Bruijn graphs constructed by the SPADES assembler (Bankevich *et al.*, 2012) to significantly improve the plasmid assembly (focusing on data generated using plasmid isolation techniques) as well as reconstruction of plasmid sequences from metagenomics datasets. Below we describe a novel plasmidSPAdes tool aimed at sequencing of plasmids from the WGS data. Recently, this problem was addressed in the case of long SMRT reads (Conlan *et al.*, 2014) but it remains open for datasets containing short Illumina reads that represent the lion’s share of bacterial sequencing projects.

We show that PLASMIDSPADES has the potential to massively increase the throughput of plasmid sequencing and to provide information about plasmids in thousands of sequenced bacterial genomes by re-assembling their genomes, identifying their plasmids, and supplementing the corresponding GenBank entries with the plasmid annotations. Such plasmid sequencing efforts are important since many questions about plasmid function and evolution remain open. For example, Anda *et al.* (2015) recently found a striking example of a bacterium (*Aureimonas sp. AU20*) that harbors the rRNA operon on a plasmid rather than on the chromosome.

Thus, re-sequencing 1000s of bacterial genomes with the goal to reassemble their plasmids will help to answer important questions about plasmid evolution. We illustrate how plasmidSPAdes contributes to plasmid discovery by analyzing *C. freundii CFNIH1* genome with well-annotated plasmids and identifying a new previously overlooked plasmid in this genome as well as discovering 7 new plasmids in ten randomly chosen bacterail datasets in the Short Reads Archive. We further provide the first analysis of accuracy of a plasmid sequencing tool across a wide variety of diverse bacterial genomes.

## 2 METHODS

### 2.1 Separating plasmids from chromosomes by read coverage

PLASMIDSPADES uses the read coverage of contigs to assist in distinguishing between plasmids and chromosomes. Illumina DNA sequencing platform typically produces reads with highly uniform coverage of the bacterial chromosomes. Fig. 1 and Table 1 illustrate that 92% (78 out of 85) of contigs greater than 10 kb in length in the assembly graph of the *E. coli* genome have coverage within 10% of the median value and 99% (84 out of 85) have coverage within 20% of the median value. The coverage of most genomes in this table is rather uniform with exception of *B. anthracis A1144, Rhodococcus J21s*, and *Thermus filiformis ATT43280*.

**Figure 1.**
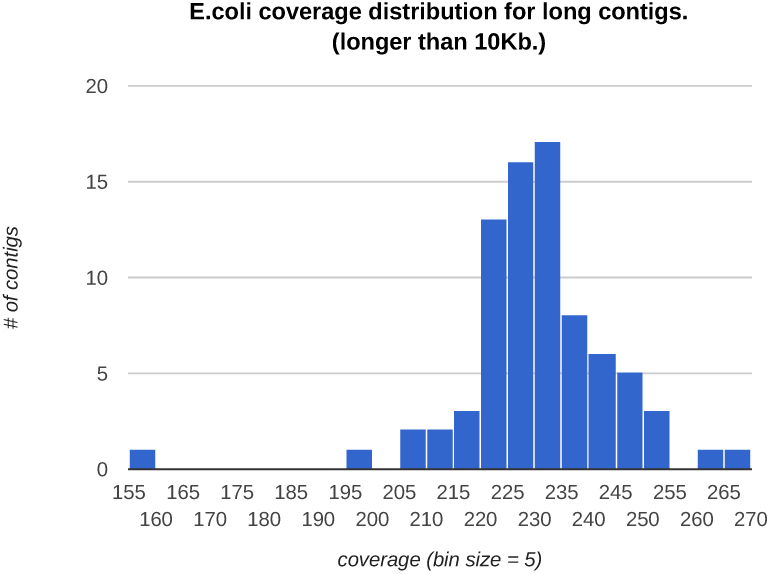
A histogram depicting the distribution of the number of long con-tigs (greater than 10 kb in length) with a given *k*-mer coverage (*k* = 55) in *E. coli* genome. The coverage of a *k*-mer in a genome is defined as the number of reads spanning this *k*-mer. The median coverage for long contigs is 231. Each bar in the histogram represents all contigs with the coverage in a bin of size 5. For example, the tallest bar in the histogram corresponds to the contigs with coverage between 230 and 235. The long contigs with minimum (160) and maximum (268) coverage have lengths 10438 bp and 13078 bp, respectively.

**Table 1.** The summary of the coverage of the contigs in various bacterial datasets. For each bacterial genome, each row shows the fraction of contigs with a coverage within 10%, 20% and 30% of the median value. For each dataset, we computed the median coverage and then counted the number of long (> 10 kb) contigs within *x*% of median coverage (for *x* = 10%, 20%, and 30%). For example, for *E. coli*, the median coverage was 216 and 85 out of 86 long contig had a coverage between within 20% of the median coverage. The table is divided into three parts corresponding to species with known plasmids (upper part), species that have no plasmids (middle part), and species for which it remains unknown whether they have plasmids. For the datasets with known plasmids (upper part of the table) we also computed the fraction of chromosomal contigs with a coverage within 10%, 20% and 30% of the median value among all *chromosomal* contigs (shown in parenthesis). The description of these datasets are provided in section 3.1 and Appendix.

| Genome | % contigs with coverage within x% of median |  |  |
| --- | --- | --- | --- |
|  | 10% | 20% | 30% |
| *B. cereus* ATCC-10987 | 66(78) | 84(100) | 84(100) |
| *R. sphaeroides* 2.4.1 | 53(73) | 68(90) | 73(97) |
| *P. stuartii* ATCC 33672 | 83(86) | 98(100) | 98(100) |
| *C. freundii* CFNIH1 | 47(55) | 86(100) | 86(100) |
| *C. callunae* DSM 20147 | 88(91) | 96(100) | 100(100) |
| *B. anthracis* A1144 | 15(14) | 23(23) | 50(50) |
| *E. coli* K12 | 92 | 99 | 99 |
| *B. cenocepacia* DDS 22E-1 | 73 | 99 | 100 |
| *Acinetobacter* UNC434CL69 | 96 | 96 | 96 |
| *Butyrivibrio* IN11a16 | 40 | 79 | 96 |
| *Lachnospiraceae* NK3A20 | 71 | 100 | 100 |
| *Luteibacter* UNC138MF | 60 | 100 | 100 |
| *Prevotellaceae* HUN156 | 100 | 100 | 100 |
| *Pseudoalteromonas* ND6B | 67 | 98 | 100 |
| *Rhodococcus* J21s | 44 | 52 | 53 |
| *Ruminococcus* YAD2003 | 64 | 100 | 100 |
| *Sphingomonas* UNC305MF | 73 | 100 | 100 |
| *Thermus filiformis* ATT43280 | 45 | 73 | 86 |

Depending on the copy number of the plasmid, its coverage can be higher or lower than the chromosome coverage. E.g., if a plasmid has a copy number 10, we expect it to have a much higher coverage than the chromosome coverage. Similarly, if a plasmid can only be found in 1/10 of the sampled cells, it will have a much lower coverage than the chromosome coverage. In order to distinguish plasmids and chromosomes by coverage, PLASMIDSPADES first estimates the chromosome coverage. The naive strategy for estimating the chromosome coverage as the average coverage over all contigs often leads to an inflated estimate because some plasmids have very large copy numbers. Since such plasmids have high coverage, the average coverage may be skewed towards the plasmid coverage.

To avoid this pitfall, PLASMIDSPADES computes the median coverage (denoted *medianCoverage*) using the assembly graph constructed by the SPADES assembler (Bankevich *et al.*, 2012). SPADES generates the assembly graph by first constructing the de Bruijn graph of all reads and further performing various *graph simplification* procedures (e.g., *bubble and tip* removals) to transform it into the assembly graph.

An edge in the assembly graph is classified as *long* if the length of the contig resulting from this edge exceeds the parameter *longEdgeLength* (the default value is 10,000 bp) and short otherwise. The median coverage is defined as the maximum coverage for which the collection of all long contigs of that coverage or greater covers at least half of the total length of the collection of all long contigs in the SPADES assembly graph. We focus on long (rather than all) contigs for two reasons. First, analyzing long contigs allows us to exclude most repeats from consideration since the longest repeats in most bacterial genomes are shorter than 10 kb (Koren *et al.*, 2013). Second, long contigs have a lower variance in their coverage than short contig. Figure 2 illustrates the larger variance in coverage for medium-sized (longer than 1 kb but shorter than 10 kb) contigs. These medium-sized contigs for the *E. coli* genome vary in coverage from 200 to 1702. For comparison, the long contigs (> 10 kb) vary in coverage from 160 to only 268.

**Figure 2.**
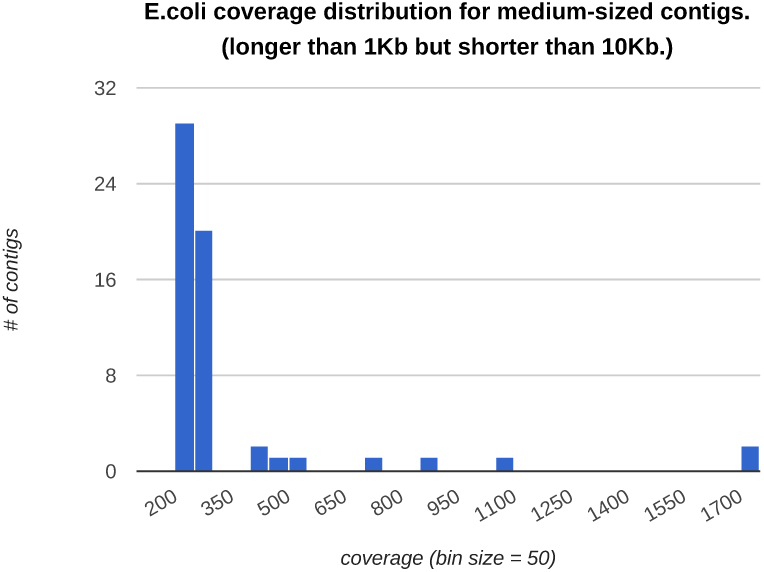
A histogram depicting the distribution of the number of medium-sized contigs (longer than 1 Kb but shorter than 10 Kb in length) with a given *k*-mer coverage (*k* = 55) in *E. coli* genome. The median coverage for medium-sized contigs is 231. Each bar in the histogram represents all contigs with the coverage in a bin of size 50. For example, the tallest bar in the histogram corresponds to the contigs with coverage between 200 and 250. The contigs with minimum (200) and maximum (1702) coverage have lengths 4122 bp and 1702 bp, respectively. Note that bars corresponding to repeats of various multiplicities are located near the projected coverage 462 (multiplicity 2), 693 (multiplicity 3), 924 (multiplicity 4), etc.

Given a parameter *maxDeviation* (the default value is 0.3), PLASMIDSPADES classifies a long edgee in the assembly graph as a *chromosomal edge* if its coverage satisfies the following condition:

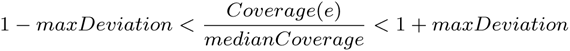

While this simple classification identifies most chromosomal contigs, it can misclassify contigs from plasmids with copy numbers close to one as being chromosomal. However, in certain cases, PLASMIDSPADES can correctly classify circular plasmids even when they have similar coverage to the chromosome. E.g., an isolated cycle in the assembly graph is classified as a putative plasmid irrespectively of its coverage.

Figure 3 shows the differences in coverage between *B. cereus* chromosome and its plasmid and illustrates the utility of using *medianCoverage* to identify chromosomal contigs. For this dataset, the long contigs can be separated with perfect sensitivity and specificity into chromosome contigs (coverage varying from 120 to 156) and plasmid contigs (coverage varying from 170 to 184) based on coverage. The *medianCoverage* (vertical green line) corresponds closely to the center of the bacterial contig distribution because bacterial genomes are typically much larger than plasmid genomes.

**Figure 3.**
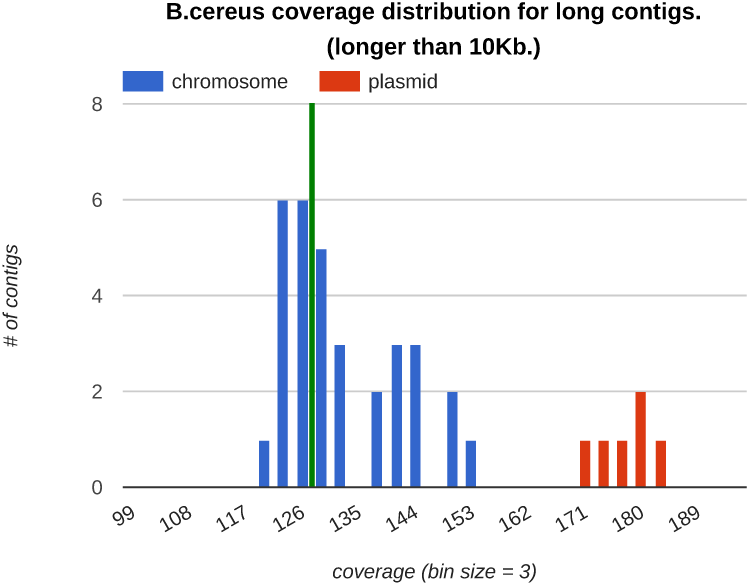
A histogram depicting the distribution of *k*-mer coverage for all long contigs in *Bacillus cereus* (*medianCoverage* = 130 marked by green line). Red bars represent contigs of chromosomal origin while blue bars represent contigs of plasmid origin. Each bar in the histogram represents all edges with the coverage in a bin of size 2.

### 2.2 plasmidSPAdes algorithm

PLASMIDSPADES utilizes SPADES for transforming the de Bruijn graph into the assembly graph (Bankevich *et al.*, 2012) and finds a subgraph of the assembly graph that we refer to as the *plasmid graph*. It further uses EXSPANDER (Prjibelski *et al.*, 2014) for repeat resolution in the plasmid graph using paired reads and generates *plasmidic contigs*.

We define the size of a connected component in the assembly graph as the sum of the lengths of the contigs resulting from its edges. An edge (*v*, *w*) in the assembly graph is called a *dead-end edge* if either the node *v* has indegree zero or the node *w* has outdegree zero (but not both). PLASMIDSPADES classifies a connected component in an assembly graph as *plasmidic* if it is composed of a single loop edge of length at least *minCirc* (default value is 1 kb) or if its size exceeds *minCompSize* (default value is 10 kb).

To transform the assembly graph into a plasmid graph, PLASMIDSPADES iteratively removes long chromosomal edges and short dead-end edges from the assembly graph. Chromosomal edges are removed because they are presumed to belong to chromosomes rather than plasmids. Dead-end edges are removed because plasmids they are not expected to generate dead-end edges.

The PLASMIDSPADES algorithm outlined below works best when the plasmids are circular and have a copy number significantly different from 1.

PLASMIDSPADES (*Reads, k, maxComponentSize*)

1. construct the de Bruijn graph using *k*-mers from *Reads* and transform it into the assembly graph.
2. compute *medianCoverage*
3. repeat
  a. remove each long chromosomal edge in the assembly graph unless it belongs to a connected component with no dead-end edges and a size less than *maxComponentSize* (default value is 150 kb)
  b. if 3.a removes at least one edge, remove all short dead-end edges and replace each non-branching path in the resulting graph with a single edge
4. remove all non-plasmidic connected components from the assembly graph to construct a plasmid graph.
5. launch EXSPANDER to perform repeat resolution on the plasmid graph
6. output all resultant plasmidic contigs and assign them to a connected component in the plasmid graph they originated from.

The *minCirc* and *minCompSize* parameters (implicit in step 4) serve an important goal of removing relatively short chromosomal contigs that evaded the step 3 of PLASMIDSPADES.

For example, error-prone reads sometimes aggregate into short paths in the assembly graph that are represented by short isolated edge with low coverage. These erroneous contigs are not removed in step 3.a because they are short and their coverage differs from the genome coverage. They are not removed in step 3.b because they are not connected to any long edges. However, they are removed in step 4. Step 6 aggregates plasmid contigs into connected components (that are expected to originate from the same plasmid) rather than outputting all plasmid contigs as a set without attempting to assign them to individual plasmids.

Ideally, PLASMIDSPADES should capture all plasmids and no chromosomal fragments in the plasmid graph (with the exception of chromosomal segments that share highly similar segments with chromosomes). However, it is not entirely true since some short segments of plasmids are sometimes missing from the plasmid graph and some short chromosomal segments are often present in the plasmid graph. Also, it is difficult to distinguish tandem repeats from plasmids by analyzing the assembly graph. Indeed, tandem repeats often form *whirls* in the assembly graph (Pevzner *et al.*, 2004) that form cycles with high coverage by reads. To distinguish plasmid from tandem repeats, one should perform the plasmid-diagnostic tests for each putative plasmid identified by PLASMIDSPADES, e.g. tests on the presence of the plasmid replication initiation protein.

## 3 RESULTS

### 3.1 Datasets

Accession numbers and links to the datasets and reference genomes are available in the Appendix:Datasets.

#### 3.1.1 Genomes with annotated plasmids

Table 2 describes six datasets that we used for benchmarking PLASMIDSPADES (see the next section for a detailed description of all columns). These datasets are composed from paired-end Illumina reads (at least 100 bp in length) from *B. cereus ATCC-10987*, *R. sphaeroides 2.4.1, P. stuartii ATCC 33672, C. freundii CFNIH1, B. cenocepacia DDS 22E-1, and C. callunae DSM 20147*. These genomes, abbreviated as *Bce*, *Rsp, Ban, Pst, Cfr, Bcen*, and *Cca*, represent well studied bacterial species with completed reference genomes and annotated plasmids. The number of plasmids of different types in these datasets varied from 0 for *Bcen* to 5 for *Rsp*. The average copy numbers varied from 1.4 for *Bce* to 4.8 for *Cca*. The lengths of plasmids varied from 4109 bp for *Cca* to 272297 bp for *Cfr*. Analysis of *B. anthracis A1144* genome is excluded from analysis in Table 2 since it has highly non-uniform coverage.

**Table 2.**
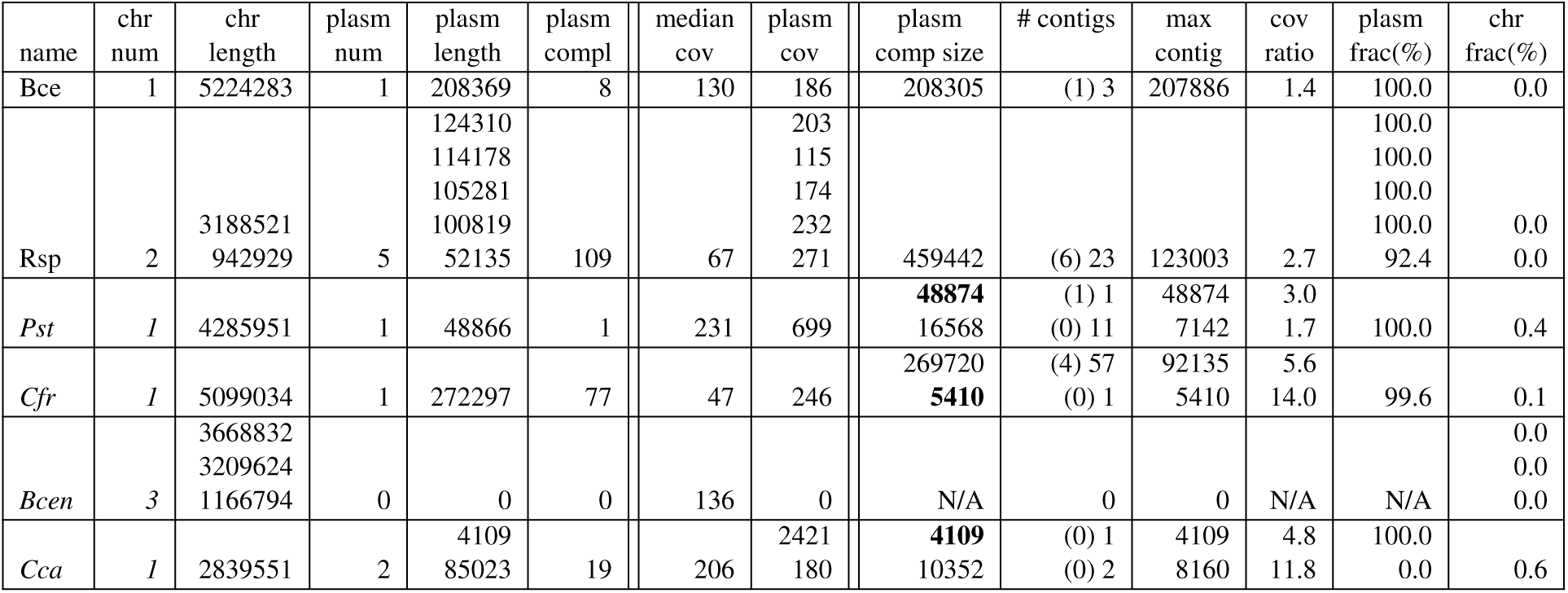
Benchmarking plasmidSPAdes on datasets with completed assemblies and annotated plasmids.

Interestingly, plasmidSPAdes assembled an additional previously unidentified short plasmid (5487 bp) in *Cfr* with high copy number (14) that is not listed in Table 2. This plasmid has a high-scoring BLAST hit to the plasmid pCAV1335-5410 in *Klebsiella oxytoca* strain CAV1335 (alignment length 4454 and percent identity 99.9).

#### 3.1.2 Genomes with unannotated plasmids

Table 3 describes ten datasets that we used for benchmarking PLASMIDSPADES for the cases when the plasmids have not been annotated yet (see the next section for a detailed description of all columns). These datasets are composed from paired-end Illumina reads (at least 100 bp in length) from *Acinetobacter sp. UNC434CL69Tsu2S25, Butyrivib-rio sp. INlla16, Lachnospiraceae bacterium NK3A20, Luteibacter sp. UNC138MFCol5.1, Prevotellaceae bacterium HUN156, Pseudoalteromonas sp. ND6B, Rhodococcus sp. J21, Ruminococcus flavefaciens YAD2003, Sphingomonas sp. UNC305MFCol5.2*, and *Thermus filiformis ATT43280*. These genomes are abbreviated as *Aci, But, Lac, Lut, Pre, Pse, Rho, Rum, Sph*, and *Tfi* in Table 3. Datasets *Aci, But, Lac, Lut, Pre* and *Rum* were downloaded from JGI read archive while datasets *Pse, Rho, Sph*, and *Tfi* were downloaded from NCBIs SRA.

**Table 3.** Benchmarking plasmidSPAdes on datasets with non-completed assemblies and lacking annotated plasmids. We boldfaced the putative plasmid composed of a single circular contig (the edge representing the contig is a loop edge).

| name | plasm comp size | # contigs | max contig | med cov | cov ratio | conf |
| --- | --- | --- | --- | --- | --- | --- |
| Aci | 66431 | (2) 20 | 32433 | 249 | 9.9 | Y |
|  | **42116** | (1) 1 | 42116 |  | 1.8 | N (phage) |
|  | 30022 | (0) 34 | 7477 |  | 6.3 | N (mobile) |
|  | **3964** | (0) 1 | 3964 |  | 15.1 | N/A |
| But | 146094 | (2) 36 | 63050 | 130 | 2.9 | Y |
|  | 34095 | (1) 1 | 34095 |  | 2.8 | N/A (plasmid) |
|  | **5609** | (0) 1 | 5609 |  | 4.1 | N/A (mobile) |
|  | **1899** | (0) 1 | 1899 |  | 3.5 | N/A (mobile) |
| Lac | **2296** | (0) 1 | 2296 | 188 | 5.9 | N/A (mobile) |
| Lut | **1445** | (0) 1 | 1445 | 227 | 6.6 | N (mobile) |
| Pre | 0 | 0 | 0 | 211 | N/A | N/A |
| Pse | 24513 | (0) 15 | 9915 | 195 | 5.1 | N |
| Rho | 830258 | (25) 120 | 112428 | 127 | 1.9 | Y |
|  | 138160 | (3) 3 | 52367 |  | 2.8 | N |
|  | **2583** | (0) 1 | 2583 |  | 7.5 | Y |
| Rum | **9168** | (0) 1 | 9168 | 157 | 3.7 | N/A (plasmid) |
| Sph | 0 | 0 | 0 | 171 | N/A | N/A |
| Tfi | 219971 | (7) 27 | 40017 | 388 | 2.3 | N |
|  | 33615 | (0) 36 | 6474 |  | 5.3 | Y |
|  | **25201** | (1) 1 | 25201 |  | 1.5 | N (rRNA) |
|  | **5150** | (0) 1 | 5150 |  | 3.7 | N/A |
|  | **1626** | (0) 1 | 1626 |  | 2.2 | N/A |

### 3.2 Benchmarking plasmidSPAdes

#### 3.2.1 Genomes with annotated plasmids

Table 2 lists the following statistics that were generated using QUAST software (Gurevich *et al.*, 2013) as well as the assembly evaluation software developed specifically for PLASMIDSPADES. The first eight columns in this table refer to the annotated chromosomes and plasmids and the remaining columns refer to the predicted plasmids.

- Species name (*name*)
- Number of chromosomes (*chr num*)
- Chromosome lengths in kb (*chr length*)
- Number of plasmids (*plasm num*).
- Plasmid lengths in bp (*plasm len*). This field lists the length of the annotated plasmid.
- Plasmid complexity reflecting the repeat content of the plasmids sequences (column *plasm compl*). To evaluate this parameter, we constructed the assembly graph from the plasmid reference sequences and counted the number of edges in this graph.
- Median coverage of the dataset (*med cov*)
- Median coverage of each annotated plasmid (*plasm cov*).
- Total length of contigs in each putative plasmid measured in bp (*plasm comp size*). We boldfaced the putative plasmid composed of a single circular contig (the edge representing the contig is a loop edge).
- Number of long contigs in each putative plasmid (shown in parenthesis) and the number of all contigs in each putative plasmid (*# contigs*)
- Longest contig in each putative plasmid (max contig)
- Coverage ratios for each putative plasmid, i.e., the coverage of each putative plasmid divided by the median coverage (*cov ratio*)
- Fraction of annotated plasmids (in percents) covered by contigs in the plasmide graph as found by QUAST (*plasm frac*). Ideally, plasmid fraction is 100%.
- Fraction of chromosome (in percents) covered by contigs in the plasmidic graph as computed by QUAST (*chr frac*). Ideally, chromosome fraction is 0%.

Appendix “Evaluating PLASMIDSPADES on annotated plasmids provides additional information about this benchmarking.

#### 3.2.2 Genomes with unannotated plasmids

Table 3 lists the following statistics that represent the PLASMIDSPADES output.

- Species name (*name*)
- Total length of contigs in each putative plasmid measured in bp ( *pl comp size*).
- Number of long contigs in each putative plasmid (shown in parenthesis) and the number of all contigs in each putative plasmid (*# contigs*)
- Longest contig in each putative plasmid (*max contig*)
- Median coverage of the dataset (*med cov*)
- Coverage ratios for each putative plasmid, i.e., the coverage of each putative plasmid divided by the median coverage (*cov ratio*)
- Confirmation status (*conf*). A “Y” indicates that the putative plasmids best blast hit to NCBI NT database was to a plasmid. A “N” indicates that the best blast hit to NT database was identified within a chromosome ( for some related species). “N/A” indicates that there are no significant matches to NCBI NT database. We further analyzed all plasmids annotated as “N” or “N/A” and, whenever possible, classified them as “N/A (plasmid)”, “N/A (phage), “N/A (rRNA)”, ‘N (phage), or “N (mobile).”

In order to validate our results, we ran a *blastn* search of longest contigs from putative plasmids constructed by PLASMIDSPADES against the NCBI database of non redundant nucleotides (NT). The best BLAST hit, defined as the hit with the lowest e-value (ties are broken by the highest bit score) for each putative plasmid component from this search can be used to identify each of the components. If the best hit for a component is to a plasmid sequence in the NT database, then we classify it as a confirmed plasmid and mark by Y in the last column of Table 3. We note that this confirmation approach is unable to confirm still unknown plasmids that have little similarity with known plasmids and so is prone to false negatives.

To further investigate the putative plasmids marked as N/A, we annotated them using RAST server (Aziz *et al.* (2008)) to check if they harbor plasmid-specific proteins. If RAST identified plasmid-specific proteins, we labeled the corresponding putative plasmidic component as “N/A (plasmid)” to emphasize that it likely represents a previously unknown plasmid. Interestingly, we found that some putative plasmids annotated as “N/A” or “N” likely represent phages (labeled as “N/A (phage)” or “N (phage).” Also, one of the putative plasmids annotated as “N/A” harbored an rRNA gene cluster (labeled as “N/A (rRNA)”).

We also conducted further analysis of putative plasmids annotated as “N” and found that many of them are formed by mobile elements that plasmidSPAdes failed to remove from the assembly graph (labeled as “N (mobile)”)‥

Appendix “Evaluating PLASMIDSPADES on unannotated plasmids” provides additional information about this benchmarking.

#### 3.2.3 Analyzing sequencing datasets with non-uniform read coverage

To investigate why some datasets (like *B. anthracis A1144* dataset) have a rather non-uniform coverage, we ordered contigs along the *B. anthracis A1144* genome and represented each contig as a bar of height equal to the coverage of this contigs. The resulting histogram (Figure 4) reveals a characteristic shape with the minimum around position 2.5 Mb. The shape of the histogram in Figure 4 is similar to the shape of the *skew diagrams* that are used for identifying the origin of replication in bacterial genomes (Compeau and Pevzner, 2015). This similarity between the histogram of coverage and the skew diagram (albeit with a few outliers) suggests that the *B. anthracis A1144* culture was sequenced in the growth phase when some cells have been replicating. The abundance of replicating cells leads to the increased coverage near the origin of replication (around position 0 Mb) and the decreased coverage near the origin of termination (around position 2.5 Mb) in *B. anthracis A1144*. See Korem *et al.* (2015) for the link between uneven coverage, rate of bacterial growth, and the origin of replication. Since the coverage of the *B. anthracis A1144* dataset is non-uniform, PLASMIDSPADES (run with the default parameter *maxDeviation* = 0.3) is unable to remove a significant fraction of chromosomal edges in the assembly graph. Since the median coverage of the *B. anthracis A1144* dataset is 130, PLASMIDSPADES only removes the edges with coverage exceeding 169 thus retaining a large number of chromosomal edges (Figure 4). While increasing the *maxDeviation* parameter to 0.4 or even higher removes nearly all chromosomal edges, it also removes some plasmidic edges ( *B. anthracis A1144* has two plasmids with coverage 135 and 183, respectively). This example illustrates additional challenges in reconstructing plasmids from sequencing datasets with highly non-uniform coverage.

**Figure 4.**
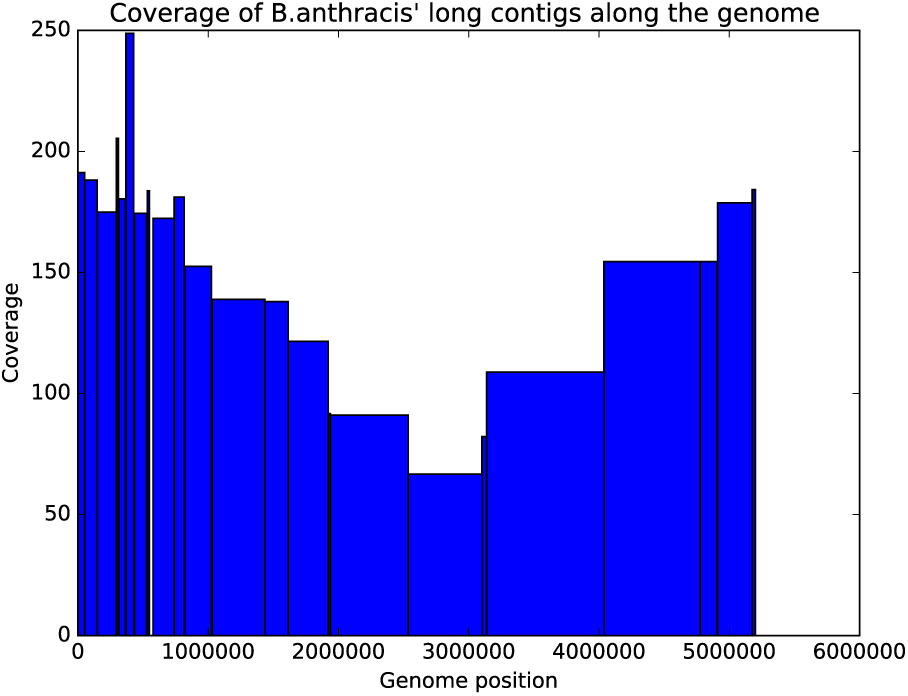
A histogram depicting the distribution of coverage of long contigs along the *B. anthracis A1144* genome.

## 4 DISCUSSION

We described a novel PLASMIDSPADES algorithm for assembling plasmids from the whole genome sequencing data. Since PLASMIDSPADES does not require any specialized sample preparation to isolate plasmids, it has a potential to increase the throughput of plasmid discovery. It thus complements the recently published approach that is mainly aimed at analyzing plasmids after plasmid isolation (Carattoli *et al.*, 2014). As Table 3 illustrates, PLASMIDSPADES identifies 7 new plasmids in 10 randomly selected SRA dataset (two of them are not similar to any previously identified plasmids). We thus expect that 1000s of new plasmids will be identified when PLASMIDSPADES is run on all SRA datasets representing bacterial and archael genomes.

PLASMIDSPADES uses coverage to remove chromosomal contigs from the assembly while retaining the plasmid contigs. We have demonstrated that in many cases it successfully removes over 99% of the chromosomal contigs while retaining over 99% of the plasmid contigs. However, PLASMIDSPADES is limited in several ways:

- In most cases, it cannot reliably identify plasmids with near chromosomal coverage.
- Since short edges in the assembly graph show a high variation in their coverage, PLASMIDSPADES cannot reliably classify short edges in the assembly graph. This makes detecting small plasmids with low copy number difficult.
- Since chromosomes are typically much longer than plasmids, if even a small portions of a chromosome is not filtered out, the small percent remaining can result in a significant number of the false positive putative plasmidic contigs caused by repetitive regions.

In the future, we plan to improve PLASMIDSPADES by adding the following features:

- When two plasmids share highly similar sequences, they may assemble into the same connected component in the assembly graph. If these plasmids have significantly different copy numbers, it may be possible to separate them from each other by analyzing their coverage using methods similar to the one developed in Rozov *et al.* (2015). E.g., PLASMIDSPADES merged five annotated plasmids in *Rsp* genomes into a single component in the plasmid graph (Table 2). However, four of them feature significantly different coverages thus enabling a possibility to identify individual plasmids from this connected component.
- Identification of plasmidic contigs can be improved by using gene prediction software, along with plasmid-diagnostic sequence features using methods similar to Lanza *et al.* (2014). E.g., because plasmids are often self-regulated, they contain certain plasmid genes to control their regulation. Similarly to PLACNET, these genes could be used as the basis for plasmid classification.

## ACKNOWLEDGMENTS

We are grateful to Anton Korobeynikov and the SPADES development team for many thoughtful discussions that helped to improve the paper.

The sequence data for *Acinetobacter sp. UNC434CL69Tsu2S25, Butyrivibrio sp. INlla16, Lachnospiraceae bacterium NK3A20*, and *Prevotellaceae bacterium HUN156* were produced by the US Department of Energy Joint Genome Institute http://www.jgi.doe.gov/ in collaboration with the user community.

